# Genetic structure of island and mainland populations of a Neotropical bumble bee species

**DOI:** 10.1101/027813

**Authors:** Flavio O. Flávio, Leandro R. Santiago, Yuri M. Mizusawa, Benjamin P. Oldroyd, Maria C. Arias

**Affiliations:** *Departamento de Genética e Biologia Evolutiva, Instituto de Biociências*, *Universidade de São Paulo, Rua do Matão 277 – sala 320, São Paulo, SP 05508-090,* Brazil; *Behaviour and Genetics of Social Insects Lab, School of Biological Sciences A12*, *University of Sydney, Sydney, NSW 2006,* Australia

**Keywords:** Bombini, *Bombus morio*, islands, microsatellites, mtDNA, population genetics

## Abstract

Biodiversity loss is a global problem and island species/populations are particularly vulnerable to such loss. Low genetic diversity is one of the factors that can lead a population to extinction. Loss of bee populations is of particular concern because of the knock-on consequences for the pollination guilds that the lost bees once serviced. Here we evaluate the genetic structure of the bumble bee *Bombus morio* populations on the mainland of South East Brazil and on nearby islands. We analyzed a total of 659 individuals from 24 populations by sequencing two mitochondrial genes (*COI* and *Cytb*) and using 14 microsatellite loci. Levels of diversity were high in most of populations and were similar on islands and the mainland. Furthermore, genetic diversity was not significantly correlated with island area, although it was lower in populations from distant islands. Our data suggest that long-term isolation on islands is not affecting the population viability of this species. This may be attributed to the high dispersal ability of *B. morio*, its capacity to suvive in urban environments, and the characteristics of the studied islands.

## Introduction

Islands often play important roles as natural laboratories for the study of ecology and evolution (MacArthur & Wilson 1967; Mayr 1967; Franks 2010). For example, the distribution of animals on islands and adjacent continents was central to the development of theories about speciation by isolation and natural selection (Darwin 1859; Wallace 1869). Currently, biodiversity loss is a global problem (Dirzo & Raven 2003; Stokstad 2006; Butchart *et al.* 2010; Bálint *et al.* 2011; Hooper *et al.* 2012; CBD 2013) and island species are of particular concern because most extinctions of mammal, bird and reptile species occurred on islands (Frankham 1997; Gaston 2009). Due to the complexity of ecological systems, the extinction of a species or population may also cause loss of important ecological interactions (Diamond 1984; Gaston 2009). Changes in predator-prey relationships, for example, can cause a cascade effect at lower trophic levels.

While humans have been the main cause of island extinctions through habitat destruction, direct predation, introduction of exotic species, and spread of disease (Frankham 1998), island species/populations may become extinct due to the combination of natural demographic, environmental and genetic factors (Shaffer 1981). Genetic diversity of island populations is expected to be low due to bottlenecks, inbreeding and genetic drift (Wright 1931; Mayr 1942; Frankham 1997). It is also expected that the size of an island, its distance from the mainland, and the time elapsed since its isolation will affect the biota’s genetic diversity (Jaenike 1973; Frankham 1997). Low genetic diversity may precipitate extinction by decreasing reproduction and survival rates, and resistance to diseases (Ayala 1965; Frankham 1998; Keller & Waller 2002; Whitehorn *et al.* 2011).

Bees are one of the most abundant and efficient pollinators and are, therefore, of particular conservation concern (Heard 1999; Cortopassi-Laurino *et al.* 2006; Steffan-Dewenter & Westphal 2008; Breeze *et al.* 2011). Absence of a single bee species can reduce the effectiveness of pollination services (Brosi & Briggs 2013) and can have knock on effects at other trophic levels (Brosi *et al.* 2007). In bees, sex is determined by zygosity at a single sex-determining locus (Cook & Crozier 1995). Females arise from diploid, fertilized eggs that are heterozygous at the sex locus, whereas males arise from unfertilized eggs. However, in small or inbred populations, diploid individuals homozygous at the sex locus are produced and are male, but are either non-viable or infertile. Thus the effects of small population size, inbreeding and low genetic diversity are generally higher in bee populations than in comparable diploid organisms (Cook & Crozier 1995), increasing the bees’ extinction proneness (Zayed & Packer 2005).

*Bombus morio* Swederus 1787 is a generalist and primitively eusocial bumble bee (Michener 2007). On average, workers are about 25 mm long, but there is wide within-colony variation (Garófalo, 1980). Its broad distribution is ill-defined but it is known from Buenos Aires (Argentina), Carabobo (Venezuela) and Lima (Peru) (Moure & Sakagami, 1962; Moure & Melo, 2012). In Brazil, it is most commonly found in areas of tropical forest and coastal vegetation (Moure & Sakagami, 1962). The intranidal population of *B. morio* consists of a queen and about 60-70 workers (Laroca, 1976; Garófalo, 1978). Like other species of this genus, they usually nest on the ground under bushes and plant debris or in cavities formed by rodents, birds and termites (Moure & Sakagami 1962; Laroca 1976; Silveira *et al.* 2002; Michener 2007). The swarming process in *B. morio* happens at least twice a year (Camillo & Garófalo, 1989). It is likely that *B. morio* has strong flight capability (Moure & Sakagami 1962), but the dispersal range of the reproductives is currently unknown.

Brazil encompasses hundreds of continental islands (previously connected to the mainland) of varying size (Ângelo 1989). Up until 17,500 years ago when sea levels were more than 100 m below their current levels, these islands were connected to the mainland (Ângelo 1989; Corrêa 1996). Isolation of these islands both from each other and the mainland occurred about 12,000 years ago (Suguio *et al.* 2005), providing a natural laboratory for studying the long-term effects of genetic isolation on bees (Rocha-Filho *et al.* 2013; Boff *et al.* 2014) and other species (Pellegrino *et al.* 2005; Grazziotin *et al.* 2006; Bell *et al.* 2012). Here we evaluate the genetic structure of Brazilian *B. morio* populations on the mainland and on islands. If the level of genetic diversity on islands is significantly lower than that observed on the mainland, then this would indicate that isolation for extended periods erodes genetic diversity in these bees, potentially leading to local extinction. If, on the other hand, the genetic diversity of island populations is similar to that found in the mainland, then this would indicate that isolation, even for millennia, has not reduced genetic diversity or population viability of these bees.

## Materials and methods

### Sampled areas

We made collections from island and mainland sites as described in Table S1. Islands ranged in size from 1.1 to 451 km^2^ and are 0.1 to 38 km from the mainland (Figure 1, Table 1). We studied 10 continental islands (previously connected to the mainland) and one sedimentary island (Ilha Comprida) which arose about 5,000 years ago (Suguio *et al.* 2003).

**Table 1.**
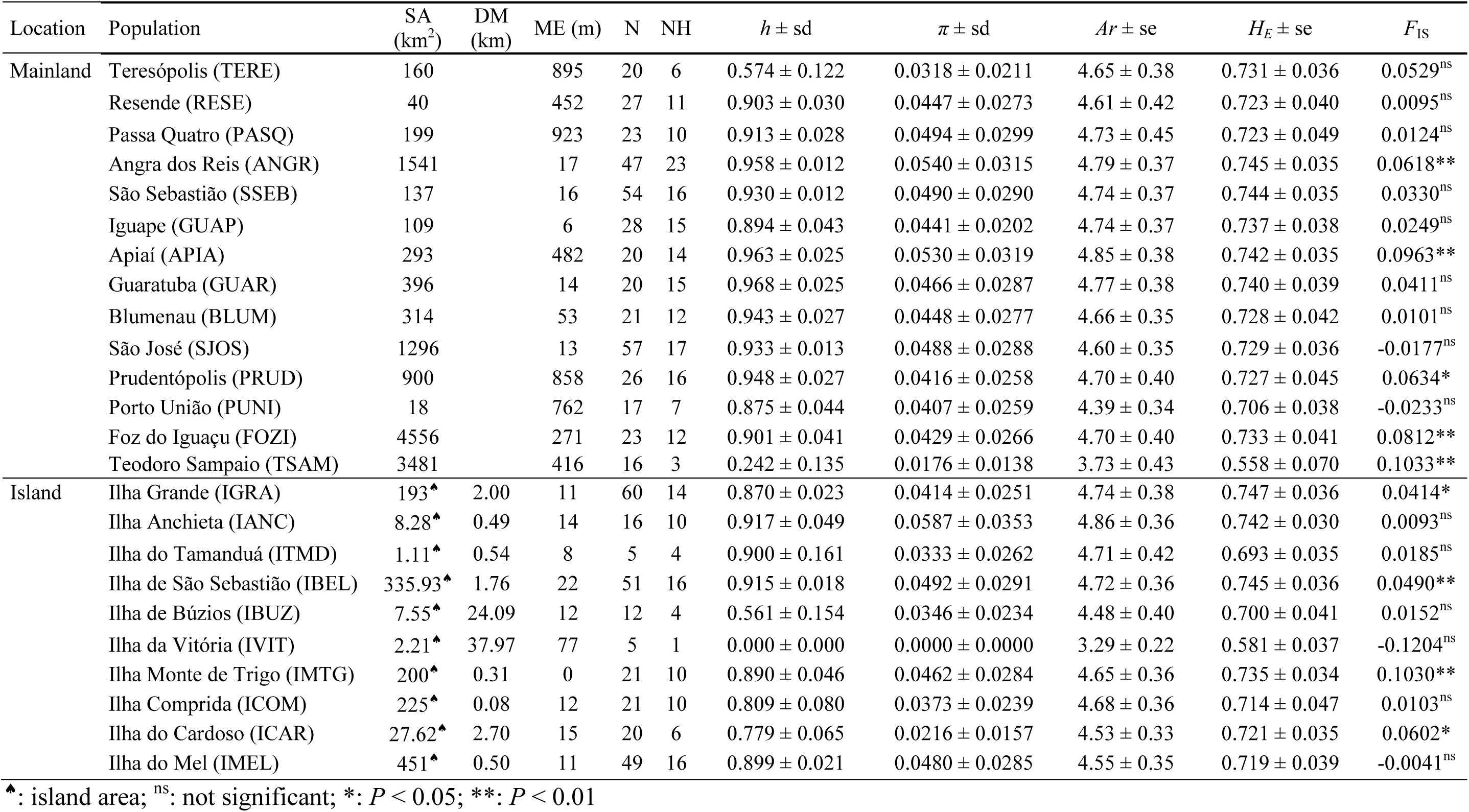
Population characteristics and genetic diversity in *Bombus morio* populations. SA: sampled area in square kilometers. ME: median elevation in meters. N: sample size. NH: number of haplotypes. *h* ± sd: haplotype diversity and standard deviation. *π* ± sd: nucleotide diversity and standard deviation. *Ar* ± se: allelic richness after rarefaction for 5 individuals and standard error. *H_E_* ± se: expected heterozigosity and standard error. *F*_IS_: inbreeding coefficient.

| Location | Population | SA<br>(km<sup>2</sup>) | DM<br>(km) | ME (m) | N | NH | $h \pm sd$ | $\pi \pm sd$ | $Ar \pm se$ | $H_E \pm se$ | $F_{IS}$ |
| --- | --- | --- | --- | --- | --- | --- | --- | --- | --- | --- | --- |
| Mainland | Teresópolis (TERE) | 160 | | 895 | 20 | 6 | $0.574 \pm 0.122$ | $0.0318 \pm 0.0211$ | $4.65 \pm 0.38$ | $0.731 \pm 0.036$ | $0.0529^{ns}$ |
| | Resende (RESE) | 40 | | 452 | 27 | 11 | $0.903 \pm 0.030$ | $0.0447 \pm 0.0273$ | $4.61 \pm 0.42$ | $0.723 \pm 0.040$ | $0.0095^{ns}$ |
| | Passa Quatro (PASQ) | 199 | | 923 | 23 | 10 | $0.913 \pm 0.028$ | $0.0494 \pm 0.0299$ | $4.73 \pm 0.45$ | $0.723 \pm 0.049$ | $0.0124^{ns}$ |
| | Angra dos Reis (ANGR) | 1541 | | 17 | 47 | 23 | $0.958 \pm 0.012$ | $0.0540 \pm 0.0315$ | $4.79 \pm 0.37$ | $0.745 \pm 0.035$ | $0.0618^{**}$ |
| | São Sebastião (SSEB) | 137 | | 16 | 54 | 16 | $0.930 \pm 0.012$ | $0.0490 \pm 0.0290$ | $4.74 \pm 0.37$ | $0.744 \pm 0.035$ | $0.0330^{ns}$ |
| | Iguape (GUAP) | 109 | | 6 | 28 | 15 | $0.894 \pm 0.043$ | $0.0441 \pm 0.0202$ | $4.74 \pm 0.37$ | $0.737 \pm 0.038$ | $0.0249^{ns}$ |
| | Apiaí (APIA) | 293 | | 482 | 20 | 14 | $0.963 \pm 0.025$ | $0.0530 \pm 0.0319$ | $4.85 \pm 0.38$ | $0.742 \pm 0.035$ | $0.0963^{**}$ |
| | Guaratuba (GUAR) | 396 | | 14 | 20 | 15 | $0.968 \pm 0.025$ | $0.0466 \pm 0.0287$ | $4.77 \pm 0.38$ | $0.740 \pm 0.039$ | $0.0411^{ns}$ |
| | Blumenau (BLUM) | 314 | | 53 | 21 | 12 | $0.943 \pm 0.027$ | $0.0448 \pm 0.0277$ | $4.66 \pm 0.35$ | $0.728 \pm 0.042$ | $0.0101^{ns}$ |
| | São José (SJOS) | 1296 | | 13 | 57 | 17 | $0.933 \pm 0.013$ | $0.0488 \pm 0.0288$ | $4.60 \pm 0.35$ | $0.729 \pm 0.036$ | $-0.0177^{ns}$ |
| | Prudentópolis (PRUD) | 900 | | 858 | 26 | 16 | $0.948 \pm 0.027$ | $0.0416 \pm 0.0258$ | $4.70 \pm 0.40$ | $0.727 \pm 0.045$ | $0.0634^{*}$ |
| | Porto União (PUNI) | 18 | | 762 | 17 | 7 | $0.875 \pm 0.044$ | $0.0407 \pm 0.0259$ | $4.39 \pm 0.34$ | $0.706 \pm 0.038$ | $-0.0233^{ns}$ |
| | Foz do Iguaçu (FOZI) | 4556 | | 271 | 23 | 12 | $0.901 \pm 0.041$ | $0.0429 \pm 0.0266$ | $4.70 \pm 0.40$ | $0.733 \pm 0.041$ | $0.0812^{**}$ |
| | Teodoro Sampaio (TSAM) | 3481 | | 416 | 16 | 3 | $0.242 \pm 0.135$ | $0.0176 \pm 0.0138$ | $3.73 \pm 0.43$ | $0.558 \pm 0.070$ | $0.1033^{**}$ |
| Island | Ilha Grande (IGRA) | 193<sup>▲</sup> | 2.00 | 11 | 60 | 14 | $0.870 \pm 0.023$ | $0.0414 \pm 0.0251$ | $4.74 \pm 0.38$ | $0.747 \pm 0.036$ | $0.0414^{*}$ |
| | Ilha Anchieta (IANC) | 8.28<sup>▲</sup> | 0.49 | 14 | 16 | 10 | $0.917 \pm 0.049$ | $0.0587 \pm 0.0353$ | $4.86 \pm 0.36$ | $0.742 \pm 0.030$ | $0.0093^{ns}$ |
| | Ilha do Tamanduá (ITMD) | 1.11<sup>▲</sup> | 0.54 | 8 | 5 | 4 | $0.900 \pm 0.161$ | $0.0333 \pm 0.0262$ | $4.71 \pm 0.42$ | $0.693 \pm 0.035$ | $0.0185^{ns}$ |
| | Ilha de São Sebastião (IBEL) | 335.93<sup>▲</sup> | 1.76 | 22 | 51 | 16 | $0.915 \pm 0.018$ | $0.0492 \pm 0.0291$ | $4.72 \pm 0.36$ | $0.745 \pm 0.036$ | $0.0490^{**}$ |
| | Ilha de Búzios (IBUZ) | 7.55<sup>▲</sup> | 24.09 | 12 | 12 | 4 | $0.561 \pm 0.154$ | $0.0346 \pm 0.0234$ | $4.48 \pm 0.40$ | $0.700 \pm 0.041$ | $0.0152^{ns}$ |
| | Ilha da Vitória (IVIT) | 2.21<sup>▲</sup> | 37.97 | 77 | 5 | 1 | $0.000 \pm 0.000$ | $0.0000 \pm 0.0000$ | $3.29 \pm 0.22$ | $0.581 \pm 0.037$ | $-0.1204^{ns}$ |
| | Ilha Monte de Trigo (IMTG) | 200<sup>▲</sup> | 0.31 | 0 | 21 | 10 | $0.890 \pm 0.046$ | $0.0462 \pm 0.0284$ | $4.65 \pm 0.36$ | $0.735 \pm 0.034$ | $0.1030^{**}$ |
| | Ilha Comprida (ICOM) | 225<sup>▲</sup> | 0.08 | 12 | 21 | 10 | $0.809 \pm 0.080$ | $0.0373 \pm 0.0239$ | $4.68 \pm 0.36$ | $0.714 \pm 0.047$ | $0.0103^{ns}$ |
| | Ilha do Cardoso (ICAR) | 27.62<sup>▲</sup> | 2.70 | 15 | 20 | 6 | $0.779 \pm 0.065$ | $0.0216 \pm 0.0157$ | $4.53 \pm 0.33$ | $0.721 \pm 0.035$ | $0.0602^{*}$ |
| | Ilha do Mel (IMEL) | 451<sup>▲</sup> | 0.50 | 11 | 49 | 16 | $0.899 \pm 0.021$ | $0.0480 \pm 0.0285$ | $4.55 \pm 0.35$ | $0.719 \pm 0.039$ | $-0.0041^{ns}$ |

<sup>▲</sup>: island area; <sup>ns</sup>: not significant; \*: $P < 0.05$ ; \*\*: $P < 0.01$

**Figure 1.**
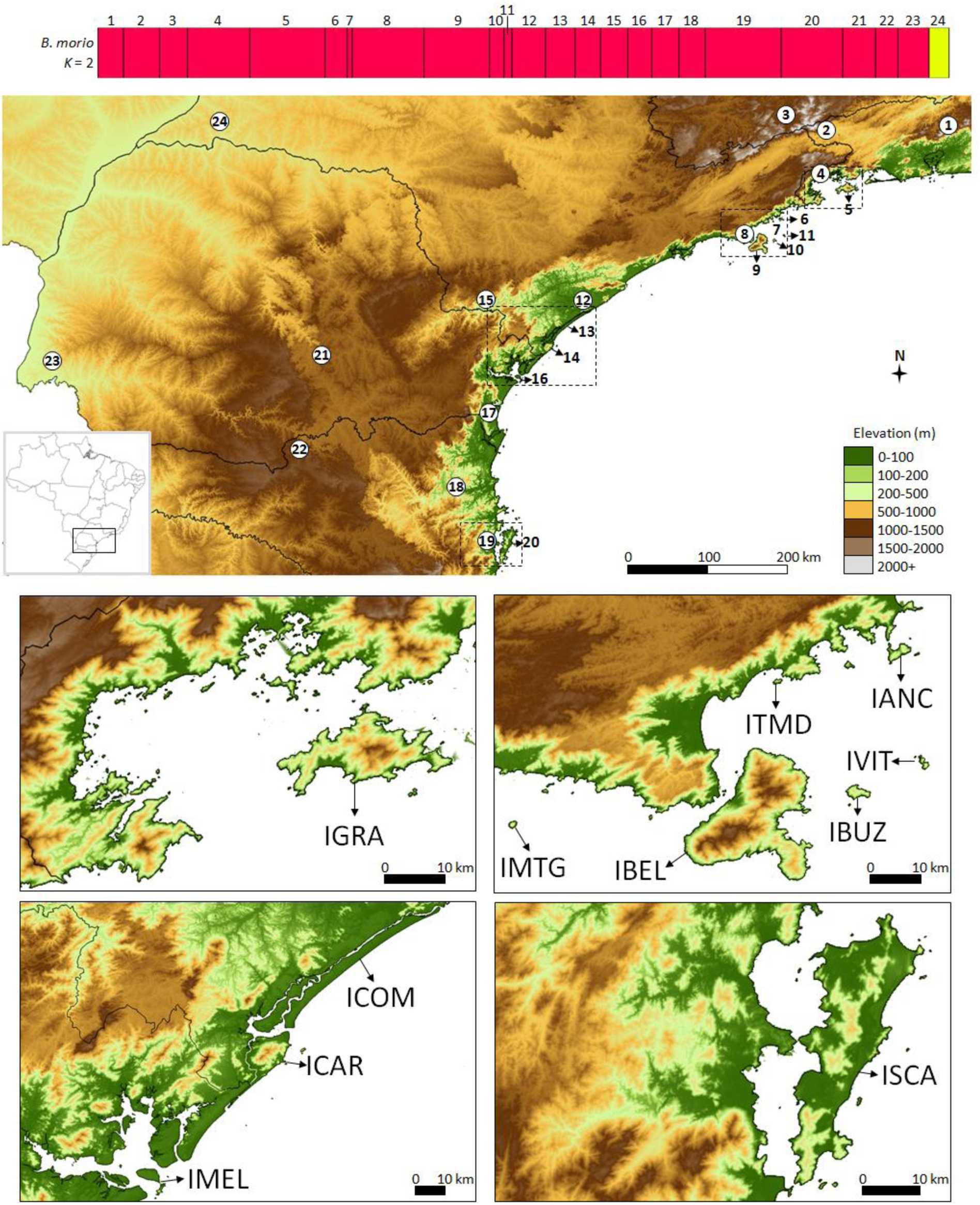
Posterior probability assignment (vertical axis) of individual genotypes (horizontal axis) for *K* = 2 in *Bombus morio* according to the program BAPS (upper panel). Below, map of the studied area with the approximate location of the sampled populations. Population names are 1: Teresópolis, 2: Resende, 3: Passa Quatro, 4: Angra dos Reis, 5: Ilha Grande, 6: Ilha Anchieta, 7: Ilha do tamanduá, 8: São Sebastião, 9: Ilha de São Sebastião, 10: Ilha de Búzios, 11: Ilha da Vitória, 12: Iguape, 13: Ilha Comprida, 14: Ilha do Cardoso, 15: Apiaí, 16: Guaratuba, 17: Ilha do Mel, 18: Blumenau, 19: São José, 20: Ilha de Santa Catarina, 21: Prudentópolis, 22: Porto União, 23: Foz do Iguaçu, and 24: Teodoro Sampaio. Detailed location of the islands visited (lower panels). IGRA: Ilha Grande; IANC: Ilha Anchieta; ITMD: Ilha do Tamanduá; IVIT: Ilha da Vitória; IBUZ: Ilha de Búzios; IBEL: Ilha de São Sebastião; IMTG: Ilha Monte de Trigo. ICOM: Ilha Comprida; ICAR: Ilha do Cardoso; IMEL: Ilha do Mel. ISCA: Ilha de Santa Catarina.

We collected 704 bees from 368 sites in Santa Catarina (SC), Paraná (PR), São Paulo (SP), Rio de Janeiro (RJ), and Minas Gerais (MG) states (Figure 1, Table S1). Samples collected in nearby cities or on the same island were grouped into the same population. In all, samples were grouped in 24 populations (Figure 1, Table 1). Bees were sampled on flowers and preserved in 96% ethanol. For DNA extraction we used one middle leg per bee. The legs were dried at room temperature for 20 min right before DNA extraction according to Walsh *et al.* (1991).

### Mitochondrial DNA sequencing

Two mitochondrial genes were partially sequenced: cytochrome c oxidase subunit 1 (*COI*) and cytochrome b (*Cytb*). Details about amplification and sequencing are given in Francisco *et al.* (2014).

### Microsatellite genotyping

We analyzed 14 microsatellite loci, 12 specific: BM1, BM3, BM4, BM5, BM7, BM9, BM10, BM11, BM12, BM13, BM17, and BM18 (Molecular Ecology Resources Primer Development Consortium *et al.* 2012) and two designed from *B. terrestris:* BT01 and BT06 (Funk *et al.* 2006). Amplification conditions of BT01 and BT06 were the same as described for BM primers, and their annealing temperatures were 48 °C and 54 °C, respectively.

Electrophoresis, visualization and genotyping were performed according to Francisco *et al.* (2011).

Micro-checker 2.2.3 (van Oosterhout *et al.* 2004) was used to verify null alleles and scoring errors. Colony 2.0.1.7 (Jones & Wang 2010) was used to verify whether individuals collected in the same plant or places nearby (less than 5 km) were related. Genepop 4.1.2 (Rousset 2008) was used to verify Hardy-Weinberg equilibrium (HWE) in populations and loci and to detect linkage disequilibrium (LD). Markov chain was set for 10,000 dememorizations, 1,000 batches and 10,000 iterations per batch. In cases of multiple comparisons *P*-values were corrected by applying Sequential Goodness of Fit test by the program sgof 7.2 (Carvajal-Rodríguez *et al.* 2009).

### Genetic diversity

Arlequin 3.5.1.3 (Excoffier & Lischer 2010) was used to calculate mitochondrial DNA (mtDNA) haplotype (*h*) and nucleotide (π) diversity. Genalex 6.5 (Peakall & Smouse 2006, 2012) was used to calculate microsatellite allelic richness and expected heterozigosity (*H_E_*). Since sample sizes were different, allelic richness was standardized (*Ar*) by rarefaction using the program hp-rare 1.0 (Kalinowski 2005). Differences in *Ar* among populations were estimated by Mann-Whitney two-tailed U Test. The inbreeding coefficient (*F*_IS_) was calculated for each population with 10,000 permutations by arlequin.

### Population differentiation

The program mega 5.2.1 (Tamura *et al.* 2011) was used to calculate the number of base substitutions per site from averaging over all sequence pairs between populations using the Kimura 2-parameter (K2p) model (Kimura 1980). Global Jost’s *D*_est_ (Jost 2008) was calculated with 9,999 permutations for mtDNA and microsatellite data by genalex. Mantel tests between genetic and geographical distances were performed with 9,999 permutations by genalex to verify isolation by distance.

Spatially clustering of individuals based on microsatellite data and the geographic coordinates was performed by Baps 6 (Corander *et al.* 2008; Cheng *et al.* 2013). The program was initially ran 5 times for each of *K* = 1 to 20 and then 10 times for each of *K* = 1 to 11. These results were used for admixture analysis with 200 iterations to estimate the admixture coefficients for the individuals, 200 simulated reference individuals per population and 20 iterations to estimate the admixture coefficients of the reference individuals.

## Results

### Sample size

We visited 11 islands and found *B. morio* on all of them except Ilha Monte de Trigo. When a bee was collected <5 km away from another bee in the collection, we determined if the two bees could possibly have come from the same colony or a different colony. If colony indicated that the two specimens could be sisters, and the two bees shared the same mtDNA haplotype we discarded one of the pair from all subsequent analyses. Overall, from the 704 bees sampled, we used 659 for population analyses (Table 1).

### MtDNA diversity

Typically, we obtained 392 bp of sequence from the *COI* gene (GenBank accession numbers KM505163-KM505866) and identified 33 haplotypes. We generated 403 bp of sequence from the *Cytb* gene had (KM505867-KM506570) and detecting 53 haplotypes. We used the concatenated sequences (795 bp) for all population analyses. All 659 sequences from the 24 populations generated 100 haplotypes (Table S2). The number of haplotypes per population ranged from 1 (Ilha da Vitória) to 23 (Angra dos Reis) (Table 1). Since *h* and *π* are correlated (*r* = 0.881, *P* < 0.001, *n* = 24) we hereafter use *π* as our measure of mtDNA diversity. High mtDNA diversity was found in all populations but Ilha da Vitória and Teodoro Sampaio (Table 1).

For mainland populations, mtDNA diversity was not significantly correlated with sampling area (*r* = −0.074, *P* = 0.731, *n* = 14) or median elevation (*r* = −0.034, *P* = 0.876, *n* = 14). For island populations, mtDNA diversity was not significantly correlated to island area (*r* = 0.475, *P* = 0.166, *n* = 10), but it was negatively correlated to the distance from the mainland (*r* = −0.748, *P* = 0.013, *n* = 10).

### Microsatellite diversity

After the Sequential Goodness of Fit correction, locus BT01 showed significant deviation from HWE in Ilha de São Sebastião (*P* = 0.004) and Prudentópolis (*P* = 0.005). Loci BM9 and BM17 showed deviation from HWE in Apiaí (*P* = 0.007) and Ilha Comprida (*P* = 0.001), respectively. Since those were occasional instances, no locus was removed from analyses. No significant LD was found between any pair of loci (all *P* > 0.05).

The number of alleles per locus ranged from four to 26, with an average of 14.1 ± 1.6 (Table S3). Mean *H_E_* was 0.75 ± 0.04. *Ar* was standardized for 5 individuals and ranged from 3.3 (Ilha da Vitória) to 4.9 (Ilha Anchieta) (Table 1). Ilha da Vitória and Teodoro Sampaio also showed the lowest *Ar* and *H_E_* values (Table 1). *Ar* was not significantly different between Ilha da Vitória and Teodoro Sampaio (*U* = 86, *P* = 0.0581), but it was between these two populations and the others (*U* < 47, *P* < 0.05). *Ar* and *H_E_* were positively correlated (*r* = 0.936, *P* < 0.001) and *Ar* will be used as indicative of microsatellite diversity hereafter.

As observed for mtDNA data, microsatellite diversity of mainland populations was not significantly correlated with sampling area (*r* = −0.171, *P* = 0.424, *n* = 14) or median elevation (*r* = −0.031, *P* = 0.884, *n* = 14). Microsatellite diversity of island populations was not significantly correlated to island area (*r* = 0.284, *P* = 0.427, *n* = 10), but it was negatively correlated to the distance from the mainland (*r* = −0.885, *P* = 0.001, *n* = 10).

Nine populations had *F*_IS_ significantly greater than zero (*P* < 0.05); five from the mainland (Angra dos Reis, Apiaí, Prudentópolis and Teodoro Sampaio) and four from islands (Ilha Grande, Ilha de São Sebastião, Ilha Comprida and Ilha do Mel). The highest *F*_IS_ (0.10) was found in Teodoro Sampaio.

### Diversity between mainland and islands

Populations were grouped according to their location: mainland or islands (Table 1). MtDNA diversity was higher in populations from the mainland (0.0435 ± 0.0263) than populations from the islands (0.0370 ± 0.0236). *Ar* was standardized for 260 individuals and populations from the mainland showed high diversity (12.65), followed by islands (11.71). Mann-Whitney two-tail U Test showed no significant differences among *Ar* values (*U* = 105, *P* = 0.748).

### MtDNA differentiation

Fourty seven of the 100 concatonated haplotypes were shared by two or more populations. We built a haplotype network where the frequency and distribution of haplotypes are shown (Figure S1). The network features a high number of interrelationships among the haplotypes and that a striking number of nucleotide substitutions separate the Teodoro Sampaio population from the others.

Global *D*_est_ was 0.344 (*P* < 0.001). The highest differentiation based on K2p was between the Teodoro Sampaio population relative to all other populations (2.106% to 2.512%) (Table 2). Teresópolis also showed substantial divergence from other populations. Mostly, however, populations were poorly differentiated. Mantel tests showed a significant positive correlation between geographic and K2p distances (*r* = 0.259, *P* = 0.046, *n* = 276).

**Table 2.** Estimates of evolutionary divergence of mitochondrial DNA sequence pairs between populations of *Bombus morio.* The number of base substitutions per site obtained from averaging over all sequence pairs between populations are shown. Analyses were conducted using the Kimura 2-parameter model (Kimura, 1980) and involved 659 nucleotide sequences. Population abbreviations as in Table 1.

|  | TERE | RESE | PASQ | ANGR | IGRA | IANC | ITMD | SSEB | IBEL | IBUZ | IVIT | GUAP | ICOM | ICAR | APIA | GUAR | IMEL | BLUM | SJOS | ISCA | PRUD | PUNI | FOZI |
| --- | --- | --- | --- | --- | --- | --- | --- | --- | --- | --- | --- | --- | --- | --- | --- | --- | --- | --- | --- | --- | --- | --- | --- |
| RESE | 0.005 |  |  |  |  |  |  |  |  |  |  |  |  |  |  |  |  |  |  |  |  |  |  |
| PASQ | 0.005 | 0.004 |  |  |  |  |  |  |  |  |  |  |  |  |  |  |  |  |  |  |  |  |  |
| ANGR | 0.005 | 0.004 | 0.004 |  |  |  |  |  |  |  |  |  |  |  |  |  |  |  |  |  |  |  |  |
| IGRA | 0.005 | 0.004 | 0.004 | 0.004 |  |  |  |  |  |  |  |  |  |  |  |  |  |  |  |  |  |  |  |
| IANC | 0.005 | 0.004 | 0.004 | 0.004 | 0.004 |  |  |  |  |  |  |  |  |  |  |  |  |  |  |  |  |  |  |
| ITMD | 0.004 | 0.003 | 0.004 | 0.004 | 0.003 | 0.004 |  |  |  |  |  |  |  |  |  |  |  |  |  |  |  |  |  |
| SSEB | 0.006 | 0.004 | 0.004 | 0.004 | 0.004 | 0.004 | 0.004 |  |  |  |  |  |  |  |  |  |  |  |  |  |  |  |  |
| IBEL | 0.006 | 0.004 | 0.004 | 0.004 | 0.004 | 0.004 | 0.004 | 0.004 |  |  |  |  |  |  |  |  |  |  |  |  |  |  |  |
| IBUZ | 0.004 | 0.003 | 0.003 | 0.004 | 0.003 | 0.004 | 0.002 | 0.004 | 0.004 |  |  |  |  |  |  |  |  |  |  |  |  |  |  |
| IVIT | 0.003 | 0.003 | 0.003 | 0.003 | 0.002 | 0.003 | 0.001 | 0.003 | 0.003 | 0.002 |  |  |  |  |  |  |  |  |  |  |  |  |  |
| GUAP | 0.005 | 0.003 | 0.004 | 0.004 | 0.003 | 0.004 | 0.003 | 0.004 | 0.004 | 0.003 | 0.003 |  |  |  |  |  |  |  |  |  |  |  |  |
| ICOM | 0.004 | 0.004 | 0.004 | 0.004 | 0.004 | 0.004 | 0.003 | 0.004 | 0.004 | 0.003 | 0.003 | 0.004 |  |  |  |  |  |  |  |  |  |  |  |
| ICAR | 0.004 | 0.004 | 0.004 | 0.004 | 0.003 | 0.004 | 0.003 | 0.004 | 0.004 | 0.003 | 0.002 | 0.003 | 0.003 |  |  |  |  |  |  |  |  |  |  |
| APIA | 0.005 | 0.004 | 0.004 | 0.004 | 0.004 | 0.004 | 0.004 | 0.004 | 0.004 | 0.004 | 0.003 | 0.004 | 0.004 | 0.004 |  |  |  |  |  |  |  |  |  |
| GUAR | 0.005 | 0.004 | 0.004 | 0.004 | 0.004 | 0.004 | 0.003 | 0.004 | 0.004 | 0.003 | 0.003 | 0.004 | 0.004 | 0.004 | 0.004 |  |  |  |  |  |  |  |  |
| IMEL | 0.004 | 0.003 | 0.003 | 0.003 | 0.003 | 0.003 | 0.002 | 0.003 | 0.003 | 0.002 | 0.001 | 0.003 | 0.003 | 0.002 | 0.003 | 0.003 |  |  |  |  |  |  |  |
| BLUM | 0.005 | 0.003 | 0.004 | 0.004 | 0.004 | 0.004 | 0.003 | 0.004 | 0.004 | 0.003 | 0.003 | 0.003 | 0.004 | 0.003 | 0.004 | 0.003 | 0.003 |  |  |  |  |  |  |
| SJOS | 0.004 | 0.004 | 0.004 | 0.004 | 0.004 | 0.004 | 0.003 | 0.004 | 0.004 | 0.003 | 0.003 | 0.004 | 0.004 | 0.003 | 0.004 | 0.004 | 0.003 | 0.004 |  |  |  |  |  |
| ISCA | 0.005 | 0.004 | 0.004 | 0.004 | 0.004 | 0.004 | 0.003 | 0.004 | 0.004 | 0.003 | 0.003 | 0.004 | 0.004 | 0.003 | 0.004 | 0.004 | 0.003 | 0.004 | 0.004 |  |  |  |  |
| PRUD | 0.005 | 0.003 | 0.004 | 0.004 | 0.004 | 0.004 | 0.004 | 0.004 | 0.004 | 0.004 | 0.003 | 0.003 | 0.004 | 0.004 | 0.004 | 0.003 | 0.003 | 0.003 | 0.004 | 0.004 |  |  |  |
| PUNI | 0.005 | 0.003 | 0.004 | 0.004 | 0.004 | 0.004 | 0.004 | 0.004 | 0.004 | 0.003 | 0.003 | 0.003 | 0.004 | 0.004 | 0.004 | 0.004 | 0.003 | 0.004 | 0.004 | 0.004 | 0.003 |  |  |
| FOZI | 0.005 | 0.003 | 0.004 | 0.004 | 0.004 | 0.004 | 0.003 | 0.004 | 0.004 | 0.003 | 0.003 | 0.003 | 0.004 | 0.003 | 0.004 | 0.003 | 0.003 | 0.003 | 0.004 | 0.004 | 0.003 | 0.003 |  |
| TSAM | 0.021 | 0.024 | 0.024 | 0.024 | 0.024 | 0.025 | 0.024 | 0.025 | 0.025 | 0.025 | 0.024 | 0.024 | 0.023 | 0.023 | 0.025 | 0.024 | 0.023 | 0.024 | 0.024 | 0.024 | 0.024 | 0.025 | 0.024 |

### Microsatellite differentiation

Global *D*_est_ was low (0.071, *P* < 0.001). Pairwise comparisons also detected low population structure, since most of *D*_est_ values were low (Table 3). Highest values were detected between the Teodoro Sampaio and Ilha da Vitória populations relative to all other populations (0.218 to 0.385). Pairwise *D*_est_ was not significantly correlated with geographic distances (*r* = 0.210, *P* = 0.070, *n* = 276). The spatial cluster approach used by baps determined *K* = 2 as the most likely optimal number of clusters (probability of 99.99%) (Figure 1).

**Table 3.** Pairwise index of differentiation (*D*_est_) from microsatellite data of *Bombus morio.* Population abbreviations as in Table 1.

|  | TERE | RESE | PASQ | ANGR | IGRA | IANC | ITMD | SSEB | IBEL | IBUZ | IVIT | GUAP | ICOM | ICAR | APIA | GUAR | IMEL | BLUM | SJOS | ISCA | PRUD | PUNI | FOZI |
| --- | --- | --- | --- | --- | --- | --- | --- | --- | --- | --- | --- | --- | --- | --- | --- | --- | --- | --- | --- | --- | --- | --- | --- |
| RESE | 0.017 |  |  |  |  |  |  |  |  |  |  |  |  |  |  |  |  |  |  |  |  |  |  |
| PASQ | 0.047 | -0.004* |  |  |  |  |  |  |  |  |  |  |  |  |  |  |  |  |  |  |  |  |  |
| ANGR | 0.052 | 0.016 | 0.021 |  |  |  |  |  |  |  |  |  |  |  |  |  |  |  |  |  |  |  |  |
| IGRA | 0.042 | 0.008* | 0.013* | 0.011 |  |  |  |  |  |  |  |  |  |  |  |  |  |  |  |  |  |  |  |
| IANC | 0.048 | 0.007* | 0.016* | 0.008* | -0.001* |  |  |  |  |  |  |  |  |  |  |  |  |  |  |  |  |  |  |
| ITMD | 0.123 | 0.093 | 0.110 | 0.024* | 0.041* | -0.026* |  |  |  |  |  |  |  |  |  |  |  |  |  |  |  |  |  |
| SSEB | 0.055 | 0.009* | 0.023 | 0.021 | 0.012 | -0.005* | 0.028* |  |  |  |  |  |  |  |  |  |  |  |  |  |  |  |  |
| IBEL | 0.048 | 0.003* | 0.024 | 0.007* | 0.009* | 0.012* | 0.032* | 0.002* |  |  |  |  |  |  |  |  |  |  |  |  |  |  |  |
| IBUZ | 0.042 | -0.011* | 0.004* | 0.009* | 0.004* | -0.017* | 0.038* | -0.011* | -0.012* |  |  |  |  |  |  |  |  |  |  |  |  |  |  |
| IVIT | 0.323 | 0.276 | 0.259 | 0.236 | 0.278 | 0.247 | 0.244 | 0.252 | 0.266 | 0.227 |  |  |  |  |  |  |  |  |  |  |  |  |  |
| GUAP | 0.058 | 0.010* | 0.026 | 0.014* | 0.006* | -0.004* | 0.027* | 0.003* | 0.005* | -0.009* | 0.296 |  |  |  |  |  |  |  |  |  |  |  |  |
| ICOM | 0.062 | 0.009* | 0.037 | 0.013* | 0.012* | -0.013* | 0.005* | -0.007* | 0.002* | -0.030* | 0.218 | -0.010* |  |  |  |  |  |  |  |  |  |  |  |
| ICAR | 0.078 | 0.015* | 0.032 | 0.014* | 0.018 | 0.001* | 0.036* | 0.002* | 0.001* | -0.013* | 0.245 | -0.006* | 0.007* |  |  |  |  |  |  |  |  |  |  |
| APIA | 0.048 | -0.006* | 0.025 | 0.024 | -0.004* | -0.011* | 0.011* | 0.005* | -0.003* | -0.016* | 0.280 | -0.015* | -0.017* | -0.005* |  |  |  |  |  |  |  |  |  |
| GUAR | 0.075 | 0.019* | 0.036 | 0.023 | 0.011* | 0.003* | 0.038* | 0.023 | 0.010* | 0.013* | 0.322 | -0.007* | 0.007* | 0.003* | -0.027* |  |  |  |  |  |  |  |  |
| IMEL | 0.062 | 0.018* | 0.049 | 0.026 | 0.015* | 0.003* | 0.043* | 0.007* | 0.013* | -0.010* | 0.299 | -0.001* | 0.002* | 0.003* | -0.012* | 0.011* |  |  |  |  |  |  |  |
| BLUM | 0.060 | 0.031 | 0.032 | 0.025 | 0.017 | 0.012* | 0.078 | 0.021 | 0.013* | 0.000* | 0.275 | 0.004* | 0.013* | 0.007* | 0.009* | -0.005* | 0.012* |  |  |  |  |  |  |
| SJOS | 0.066 | 0.027 | 0.058 | 0.034 | 0.026 | 0.023 | 0.062 | 0.017 | 0.016 | 0.014* | 0.283 | 0.012* | 0.014* | 0.002* | 0.011* | 0.020 | 0.007* | 0.000* |  |  |  |  |  |
| ISCA | 0.098 | 0.050 | 0.078 | 0.053 | 0.044 | 0.043 | 0.081 | 0.026 | 0.025 | 0.021* | 0.286 | 0.030 | 0.029 | 0.007* | 0.030 | 0.034 | 0.012* | 0.002* | -0.008* |  |  |  |  |
| PRUD | 0.060 | 0.030 | 0.018* | 0.014* | 0.030 | 0.037 | 0.073 | 0.025 | 0.016 | -0.007* | 0.293 | 0.019 | 0.023* | 0.022 | 0.017* | 0.021* | 0.045 | 0.033 | 0.048 | 0.057 |  |  |  |
| PUNI | 0.076 | 0.038 | 0.030 | 0.023 | 0.039 | 0.013* | 0.052* | 0.018* | 0.034 | 0.030* | 0.255 | 0.015* | 0.018* | 0.019* | 0.038 | 0.038 | 0.048 | 0.056 | 0.055 | 0.077 | 0.043 |  |  |
| FOZI | 0.052 | 0.001* | 0.023 | 0.015* | 0.025 | 0.016* | 0.063 | 0.016* | 0.007* | -0.010* | 0.311 | 0.002* | 0.005* | 0.023 | -0.001* | 0.003* | 0.019* | 0.034 | 0.048 | 0.055 | -0.009* | 0.024* |  |
| TSAM | 0.327 | 0.261 | 0.308 | 0.231 | 0.276 | 0.240 | 0.252 | 0.286 | 0.264 | 0.262 | 0.385 | 0.255 | 0.238 | 0.255 | 0.244 | 0.264 | 0.267 | 0.301 | 0.306 | 0.309 | 0.249 | 0.230 | 0.245 |

Colours highlight $D_{\text{est}}$ values. Green: $D_{\text{est}} < 0.05$ ; yellow: $0.05 < D_{\text{est}} < 0.15$ ; orange: $0.15 < D_{\text{est}} < 0.25$ ; red: $D_{\text{est}} > 0.25$ ; \* $P > 0.05$ ; values without asterisk $P < 0.05$ .

## Discussion

The genetic diversity of *B. morio* populations is similar on mainland and island sites. Furthermore, genetic diversity is not significantly correlated with island area, although it is lower in populations from islands that are more distant from the mainlad. It is noteworthy that *B. morio* shows limited genetic divergence between island and mainland populations and among most of the mainland sampling sites. We suggest that the dispersal ability of *B. morio* combined with its capacity to live in urban environments, and the characteristics of the studied islands explain the genetic structure of the Brazilian populations.

The dispersal of *B. morio* is intimately related to its nesting and reproductive behavior. Colony reproduction in *Bombus* begins when a young queen leaves the mother nest alone and is fertilized by a male. Males and queens may have multiple partners, though this is rare for queens (Garófalo *et al.* 1986; Estoup *et al.* 1995). The mated queen begins to look for a suitable place to build the new nest. Once a nest site is found, the queen starts oviposition and performs all activities, such as foraging, cell provisioning and feeding the larvae (Garófalo 1979). When workers emerge, labor division is set (Michener 2007) and the queen never leaves the nest again (Laroca 1976). The lack of dependence on the mother nest means that daughter colonies can be established a considerable distance from the natal nest. In Europe, *Bombus* queens have been observed several kilometers off shore over water (Macfarlane & Gurr 1995; Widmer *et al.* 1998; Darvill *et al.* 2010). In New Zealand, queens of *B. terrestris* colonized islands up to 30 km from the mainland (Macfarlane & Gurr 1995).

Our data suggest that both female and male *B. morio* have high dispersal abilities, although males have higher. Some population structure and isolation by distance were detected in the mitochondrial analyses, but our microsatellite data showed negligible genetic structure even over distances exceeding 1,000 km. The homogeneity of *Bombus* populations has also been observed in European and North American populations (Estoup *et al.* 1996; Ellis *et al.* 2006; Lozier & Cameron 2009; Lozier *et al.* 2011). In *B. terrestris*, males can fly up to 10 km, including over water (Kraus *et al.* 2009). Most likely, long distance dispersal allows *B. morio* to minimize the effects of isolation on islands.

*Bombus morio* was easily found at all mainland locations, even in urban environments. *Bombus* can commonly be seen visiting flowers along roadsides (Lozier *et al.* 2011), including *B. morio* (personal observations). In Europe and North America several bumble bee species are common in urban environments because the gardens and parks provide a diversity and abundance of flowers throughout the breeding season (Chapman *et al.* 2003; Goulson 2010; Lozier *et al.* 2011). Similarly, *B. morio* thrives in Brazilian urban environments, and this ecological capacity no doubt contributes to its dispersal.

The vast majority of island and mainland populations of *B. morio* have moderate/high nuclear genetic diversity. In contrast, studies of other *Bombus* species on islands have found high differentiation and low genetic diversity (Estoup *et al.* 1996; Widmer *et al.* 1998; Shao *et al.* 2004; Darvill *et al.* 2006, 2010; Schmid-Hempel *et al.* 2007; Goulson *et al.* 2011; Lye *et al.* 2011; Lozier *et al.* 2011; Moreira *et al.* 2015). This discrepancy may be due to the fact the islands studied here are closer to the mainland than in the other studies. When we visited the most isolated islands we found low genetic diversity (Ilha da Vitória) or absence of bumble bees (Ilha Monte de Trigo).

The failure in collecting *B. morio* on Ilha Monte de Trigo may be due to any of the following: insufficient collection effort, ancestral absence from the island when it was isolated, or its extinction after isolation. Our collection effort was eight hours, so it is not possible to assert that the species does not occur on this island, although this amount of time was sufficient to find *B. morio* on all other islands. Its distance from the mainland, 10.2 km, may prevent queen (re)colonization. Nonetheless, *B. morio* is present on Ilha da Vitória, a more distant island (38 km from the mainland and 11 km from the nearest island). We only found five bees on Ilha da Vitória and its population showed low genetic diversity, suggesting that the Ilha da Vitória population is small and may be threatened. Interestingly, the population of the orchid bee *Euglossa cordata*, which like *B. morio*, has good dispersal abilities, from Ilha da Vitória has low genetic diversity and is strongly differentiated from adjacent populations, both on the nearby islands Ilha de Búzios and Ilha de São Sebastião and from those on the mainland (Boff *et al.* 2014).

*Bombus morio* mitochondrial genetic diversity is high, except in Ilha da Vitória and Teodoro Sampaio. Many populations had *h* > 0.9. In fact, bees collected off the same flower or plant often had different haplotypes. Chapman *et al.* (2003) also observed that both *B. terrestris* and *B. pascuorum* workers visiting the same plant are often from different colonies. This high genetic diversity also suggests that populations of *B. morio* did not experience genetic bottlenecks during the Pleistocene. Indeed, *Bombus* species have some characteristics as robustness, hairiness and thermoregulatory adaptations that allow them to survival in temperate and cold regions (Hines 2008).

For both markers, the island populations and its nearby mainland populations are undifferentiated, most likely because of frequent migration. For example, the Ilha da Vitória population is not differentiated from Ilha de Búzios, the nearest island, with respect to mitochondrial haplotypes. The single haplotype found on Ilha da Vitória (H3) is the most common on Ilha de Búzios, and is also found on all other populations studied, except Teodoro Sampaio. Although it is possible that this haplotype is a relict of the ancestral population formed at the time of isolation, it is more likely that it is a result of a more recent colonization by queens from Ilha de Búzios.

Inbreeding is not a current concern for the island populations we studied. Although *F*_IS_ was significantly different from zero on three island populations, *F*_IS_ was high only on Ilha Comprida. However, this population has high genetic diversity and is not genetically isolated, so the high *F*_IS_ is likely to be eroded with time.

Both markers indicated low genetic diversity in the Teodoro Sampaio population and high differentiation between Teodoro Sampaio and the other populations. Genetic drift may be the primary driver of this result. The Teodoro Sampaio population did not share haplotypes with any other population, whereas other populations all shared at least one haplotype. In addition, haplotypes are very similar, being distinguished by only one nucleotide in most cases. The percentage of variable sites between Teodoro Sampaio and other populations (from 2.065 to 2.454%) was higher than that seen among all other populations, whose maximum value was 0.554% (São Sebastião × Teresópolis). The significant genetic divergence of the Teodoro Sampaio population from all others suggests that this population is a subspecies of *B. morio*.

To our knowledge, this is the first comparative study of the genetic architecture of mainland and island populations of a Neotropical bumble bee species. Our study shows that even in a highly fragmented landscape *B. morio* survives in urban environments and enjoys a high level of genetic diversity. This suggests *B. morio* populations are self-sustaining, and that this species will remain as an important pollinator in Brazil.

## Acknowledgments

We are grateful to Paulo Henrique P. Gonçalves for his help with the sampling and to Susy Coelho and Julie Lim for technical assistance. We thank Adílson de Godoy, Carlos Chociai, Flávio Haupenthal, Geraldo Moretto, Marcos Wasilewski, Marcos Antonio, Renato Marques, José Moisés, André Trindade, Teófilo, Eduardo da Silva, Guaraci Cordeiro, Marcos Fujimoto, PC Fernandes, Samuel Boff, Thaiomara Alves, the managers and the staff of the Parks, the residents of Ilha da Vitória, Ilha de Búzios and Ilha Monte de Trigo, and countless people who assisted us in the fieldwork. We thank Dr. Jeffrey Lozier for comments on an early version of this manuscript. For permits, we thank Instituto Brasileiro do Meio Ambiente e dos Recursos Naturais Renováveis (IBAMA) and Instituto Chico Mendes de Conservação da Biodiversidade (ICMBio) (18457-1), Instituto Florestal do estado de São Paulo (260108 - 000.000.002.517/0 2008), Instituto Ambiental do estado do Paraná (128/09) and Instituto Estadual do Ambiente do Rio de Janeiro (E-07/300.011/0). This work was supported by Fundação de Amparo à Pesquisa do Estado de São Paulo (04/15801-0; 08/07417-6; 08/08546–4; 10/18716-4; 10/50597-5) and Australian Research Council. This work was developed in the Research Center on Biodiversity and Computing (BioComp) of the Universidade de São Paulo (USP), supported by the USP Provost’s Office for Research.

